# Spatio-temporal feature based deep neural network for cell lineage analysis in microscopy images

**DOI:** 10.1101/2021.08.28.457873

**Authors:** Siteng Chen, Andrew L. Paek, Kathleen A. Lasick, Suvithanandhini Loganathan, Janet Roveda, Ao Li

## Abstract

**Background:** Time-lapse microscopy has been widely used in biomedical experiments because it can visualize the molecular activities of living cells in real time. However, biomedical researchers are still conducting cell lineage analysis manually. Developing automatic lineage tracing algorithms is a challenging task. In the past two decades, deep neural networks (DNNs) became have shown outstanding performance on computer vision tasks. They can learn complex visual features, capture long-range temporal dependencies, and have the potential to be used for automatic cell lineage analysis.

**Methods:** In this study, we propose a multi-task spatio-temporal feature based deep neural network for cell lineages analysis (Cell-STN). The Cell-STN extracts spatio-temporal features from microscopy image sequences by leveraging our convolutional long short-term memory based core block. And the proposed Cell-STN utilized a task specific network to predict the cell location, the mitosis event, and the apoptosis event in a multi-task manner.

**Results:** We evaluated the Cell-STN on three in-house datasets (MCF7, U2OS, and HCT116) and one public dataset (Fluo-N2DL-HeLa). For cell tracking, we used peak-wise precision, track-wise precision, end-peak precision, and spatial distance as metrics. The overall results showed the Cell-STN models outperform other state-of-the-art cell trackers. For mitosis and apoptosis tasks, we used accuracy, F1-score, temporal distance, and spatial distance as metrics. The Cell-STN models achieved the highest performance on all datasets.

**Conclusion:** This study presented a novel DNNs approach for cell lineage analysis in microscopy images. The Cell-STN showed outstanding performance on the four datasets. Additionally, the Cell-STN required minimal training data and can be adapted to new biological event detection tasks by appending task-specific layers. This algorithm has the potential to be used in real-world biomedical research.

## Introduction

Tracking cell lineages in time-lapse microscopy images has had a major impact on our understanding of cell biology. For example, lineage tracking experiments have revealed that cell division is coupled to the circadian clock (Sandler et al., 2015) and that circadian coupling influences the fate of cancer cells during chemotherapy treatment (Chakrabarti et al., 2018). Time-lapse microscopy is also widely used to visualize the dynamics of protein activity in living cells over time (Purvis & Lahav, 2013) and has revealed these dynamics control cell division (Albeck, Mills & Brugge, 2013), differentiation (Kobayashi et al., 2009) and death (Paek et al., 2016). Yet researchers are still analyzing cell lineages manually, which is time consuming and prohibitive for large scale studies. Thus, developing automatic algorithms for tracking cells and identifying mitotic and apoptotic events will likely have a large impact on biomedical research.

Several automatic algorithms have been developed for cell lineages analysis. For example, Tian et al. introduced the EllipTrack algorithm that used a global track-linking algorithm and a local track-correction module for cell lineage analysis (Tian, Yang & Spencer, 2020). Magnusson et al. also introduced a global track linking algorithm (BaxterAlgorithm) that linked cell outlines generated by a segmentation algorithm into tracks which used Viterbi algorithm to find the tracks (Viterbi, 1967; Magnusson et al., 2015). These above two methods achieved good performance in the lineage analysis tasks. However, the training procedure of the above two algorithms required extremely accurate mask label that is expensive in terms of both time and cost in biomedical experiments. To obtain the additional labels for algorithm training, human experts have to tune a number of parameters to annotate the spatial location as well as the exact time point for each cell and biological events throughout the vision score from hundreds of video frames. In order to detect the mitosis event and apoptosis event, several works have also been proposed for these two importance sub-tasks of lineage analysis. Some works, such as the Hidden Conditional Random Fields (Liu, Li & Kanade, 2010), 3D Convolutional Neural Networks (Nie et al., 2016), used handcrafted features to model the temporal dynamics of mitotic events. As a result, the manual preprocess needs careful selection of hyper-parameters with prior domain knowledge. The pixel-level cell segmentation methods (Li et al., 2018, 2019) and fully convolutional networks (Ronneberger, Fischer & Brox, 2015; Zhou, Mao & Yi, 2017) were used for detecting mitosis events. However, these latter two methods lack the ability to learn temporal features. Approaches with little intervention are needed to avoid this time-consuming procedure.

Deep neural networks (DNNs) are a solution for automatic cell lineage analysis. In the past two decades, the DNNs have been increasingly utilized to analyze biomedical data. The DNNs can directly learn features from raw input and discover unrecognized patterns in high-dimensional data (Lipton et al., 2015). Each layer of a DNN includes multiple filters that are designed to extract features at different levels. In a classification task, higher level layers amplify aspects of the inputs that are important for discrimination and suppress irrelevant variations (LeCun, Bengio & Hinton, 2015). Therefore, this architecture of DNNs is ideal for analyses of complex nonlinear, multidimensional biomedical data.

In this work, we propose a DNNs algorithm that can automatically track cells, detect mitotic event, and apoptosis event. The proposed multi-task spatio-temporal feature based deep neural network for cell lineage analysis (Cell-STN) includes two parts. We applied a shared spatio-temporal backbone network to extract spatio-temporal features of the target cell and a three task specific networks to predict the cell location, the mitosis event, and the apoptosis event. The models were trained and tested on three in-house dataset and one public dataset. Importantly, the training process of Cell-STN is much simpler than previous algorithms. Biomedical researchers do not need to make additional annotations (e.g., accurate mask-level labels) to train the Cell-STN models. Therefore, the Cell-STN has potential for widespread use in real-world biomedical research.

## Materials & Methods

### Datasets

We conduct experiments on four datasets, including three in-house datasets collected at the University of Arizona, and one public dataset, Fluo-N2DL-HeLa, from the Cell Tracking Challenge (Maška et al., 2014; Ulman et al., 2017). All in-house datasets contained three channels: the CFP channel captured the nucleus information indicated by H2B protein; the bright-field channel captured phase-contrast images; the YFP channel was used for biomarker p53 in the HCT116 and U2OS data sets, and Foxo1 in the MCF7 data set. An example of the in-house dataset is shown in Figure S1. The details of these datasets are described as following:

#### U2OS

The U2OS dataset included 30 time-lapse microscopy movies (Paek et al., 2016). This study was performed on human U2OS bone osteosarcoma cells in 4 biological experiments. Each movie included 301 frames with a time step of 30 minutes. In the experiments, a drug addition (doxorubicin) was added between frames 148 and 149. The doxorubicin caused abnormal cell morphology and drastic changes to the image background. In each movie, 10 cells were annotated by experimentalists using morphology. We randomly split the 4 movies to training set (n=3) and testing set (n=1). We further split the training set to a training set (n=2) and a validation set (n=1) to train Cell-STN models.

#### HCT116

The HCT116 dataset consisted of 4 time-lapse microscopy movies, performed on human HCT116 colon cancer cells. In each experiment, we captured 243 images with a time step of 30 minutes. Each image was captured in 1024 × 1024 pixels and encoded in the 16-bit Tagged Image File Format. A drug addition, (cisplatin) was added between frames 99 and 100. Human experts tracked 387 cells in this experiment. The HCT116 dataset was the most challenging dataset in this study. Compared to the MCF7 and U2OS cells, the HCT116 cells were very likely to jump to a different region during division. The cisplatin also caused rapid cell migration, dense cell distribution, and strong overlap. Additionally, the objective used for this dataset has lower quality than the one for the MCF7 and U2OS dataset, which lead to blurry images. We randomly split the 4 movies to training set (n=3) and testing set (n=1). We further split the training set to a training set (n=2) and a validation set (n=1) to train Cell-STN models.

#### MCF7

The MCF7 dataset included 50 time-lapse microscopy movies. In each movie, the frames were recorded every 20 minutes for 145 times. H_2_O_2_, which induced cell death, was added between frame 3 and 4 of each movie. Each frame had a size of 1024 × 1024 pixels. The MCF7 cells moved slower than other cells in this study. Human experts labeled 500 cells for lineage analysis. In the experiments, because of different study purpose, human experts only marked the frames when cell division began and ended. And only one of the two daughter cells was tracked and labeled after division. In this study, we selected the middle point of the division period as the ground truth annotation to train Cell-STN models. We randomly split the 30 movies to a training set (n=27) and a testing set (n=3). We further split the training set to a training set (n=24) and a validation set (n=3) to train Cell-STN models.

#### HeLa

The Fluo-N2DL-HeLa dataset contained 2 microscopy movies (Maška et al., 2014; Ulman et al., 2017). In each movie, HeLa cells expressing H2B-GFP were recorded every 30 minutes for 92 frames. The moving speed of HeLa cells was similar to MCF7 cells. The apoptosis events were rare in the HeLa dataset. Because of the limit number of independent movies, we applied a 2-fold cross-validation with the sample size of one biological experiment to train and test the Cell-STN models

In this study, we split the time-lapse movies into smaller subsequences of 10 frames with 256-by-256 pixels using a moving window method with spatial steps of 128 pixels and temporal steps of 5 frames to reduce the large memory footprint of the data. We also used data augmentation technology to reduce the likelihood of overfitting during training by flipping each subsequence horizontally and vertically and rotate by 90, 180, and 270 degrees (Kumar et al., 2017).

### Model Development

In cell lineage analysis, given a set of *N* adjacent frames *I* = {*i*_1_, *i*_2_, ⋯, *i*_*N*_}, where *i*_*t*_ ∈ [0,1]^*X*×*Y*^ represents a frame of size *X* × *Y* at frame *t*, and the initial central coordinates 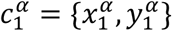 for the target cell *α*, we defined the lineage matrix *L*^*α*^ = {*C*^*α*^, *M*^*α*^, *A*^*α*^} for a target cell, where the coordinate set 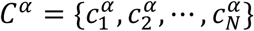, mitosis set 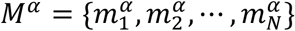, and apoptosis set 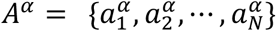. The 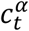 represented the spatial location of the target cell *α* in frame *i*_*t*_. If the target cell split into two daughter cells in *i_t_*, 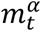 equaled 1 (presence of mitosis event); otherwise, it equaled 0. Similarly, we used 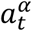 to denote the presence of apoptosis event in frame *i*_*t*_. Therefore, the problem was to solve the probability of biological events of cell *α* at each frame, given the frames *I* and the initial point 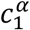 that can be expressed as 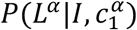.

In this study, we represented the coordinates of the target cell and biological events in spatial space: the coordinates maps 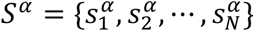 is the spatial representative of 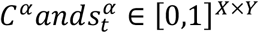, where the 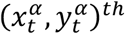 entry *s*_*t*_ equals 1; otherwise equals 0. We denoted the biological events in spatial space as the mitosis maps (Eq. 1) and apoptosis maps (Eq. 2), where ⊙ denotes the Hadamard operator. As a result, the original problem became the following three tasks, 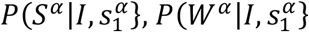, *and* 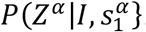.

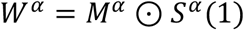

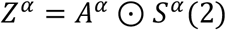

Acquiring the spatio-temporal features of the target cell, denoted as *ST*^*α*^, was the key to solve this problem. In our approach, the inputs of the Cell-STN were a sequence of cell images and the initial spatial information (the coordinates map) of the target cell. We modeled the cellular dynamics relevant to the target cell by combining the cellular dynamics for all objects and the predicted trajectory for the specific target cell. We used the observed frames *I* to learn the cell behaviors and appearance for entire population from frames 1 to *N*, and the initial coordinates map 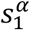 to obtain the target cell deep latent space from frames 1 to *N*.

To solve this problem, we modeled the dynamics of cellular activities, *P*(*ST*|*I*}, with *ST* being the latent state that captured the complete dynamics of cell population and predicted the latent state *Q*^*α*^ specific to the target cell *α* with the probability 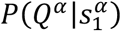. Then we constructed spatio-temporal features for the target in deep latent space by computing the probability *P*(*ST*^*α*^|*ST,Q*^*α*^). Without loss of generality, the problem was formulated with the joint probability density (Eq. 3). Consequently, we can formulate the cell tracking, mitosis detection, and apoptosis detection tasks as Eq. 4 – Eq. 6, respectively.

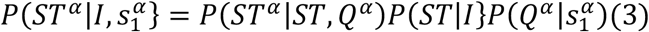

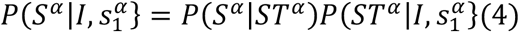

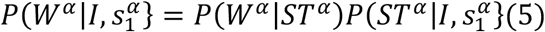

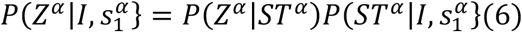

We observed the term 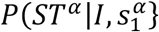 exists in Eq. 3-Eq. 6. Hence, we proposed a multi-task spatio-temporal feature based deep neural network with a shared spatio-temporal Core network to learn latent space 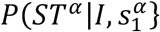, and a three task specific networks for cell tracking *P*(*S*^*α*^|*ST*^*α*^), mitosis detection *P*(*W*^*α*^|*ST*^*α*^), and apoptosis detection *P*(*Z*^*α*^|*ST*^*α*^).

### Spatio-temporal Core Network

The proposed spatio-temporal core network consisted of three branches (E, P, A). This network is depicted in Figure 1. The encoding branch (E) learnt the movements and morphology of the cell population in frames. The prediction branch (P) projected the initial coordinates map to a deep latent space and predicted the position of the target cell at a time point. The attention branch (A) combined the outputs of E and P to construct the target specific spatio-temporal features.

**Figure 1.**
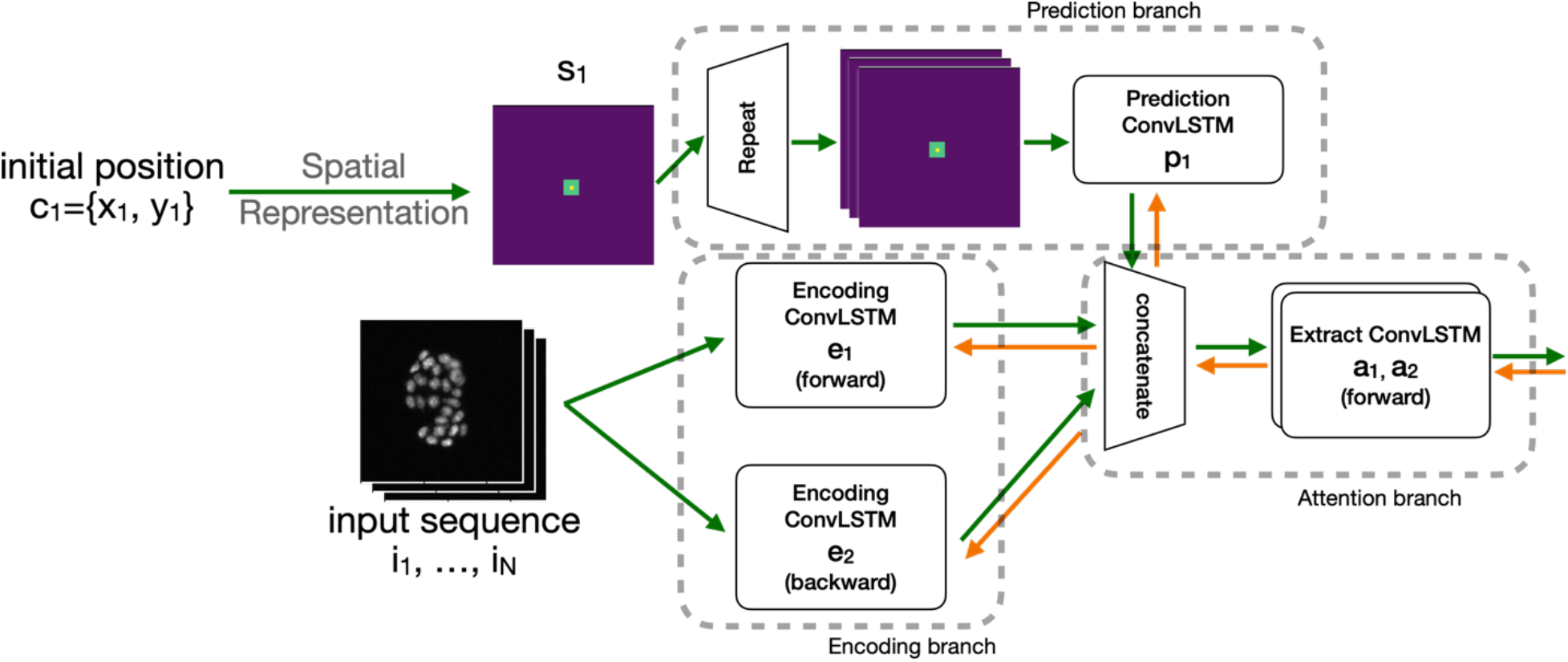
The architecture of the spatio-temporal core network. It consists of three branches: encoding branch (E) for modeling morphology and activities of the cell population, prediction branch (P) for approximating the trajectory of the target cell, and attention branch (A) for fusing the population information and individual latent state. The green arrow indicates the inference paths. The orange arrow indicates the backpropagation paths.

In the encoding branch, we apply bidirectional convolutional long-short term memory (ConvLSTM) to screen the input imaging sequence in forward and backward directions to learn the probability function *P*(*ST*|*I*). The long-short term memory (LSTM), one of the variants of RNN, was originally proposed by Hochreiter and Schmidhuber (Hochreiter & Schmidhuber, 1997). The ConvLSTM is a combination of convolutional neural network and LSTM that can preserve spatial and temporal information to model long-range dependencies and encode the spatio-temporal information of image sequences (Shi et al., 2015). The details of LSTM and ConvLSTM are described in Supplementary. Biological events, such as mitosis events, occurred in multiple phases. And each phase had different characteristics in morphology. The information of the later phases contributed to the decision-making process of a previous event. Therefore, the ConvLSTMs with two directions were used to ensure that both the past and future dynamics in the cell population are captured. In this study, the two ConvLSTMs extracted visual features and temporal patterns of the cell population. The outputs of the two ConvLSTM layers were denoted as *ST*_*forward*_ and *ST*_*backward*_, respectively.

The prediction branch was used to expand the two-dimensional initial coordinates map 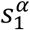 into a sequence of hidden states *Q*^*α*^, expressed as 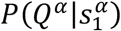,which approximated the trajectory of the target cell. The initial coordinates 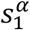 was repeated *N* times. And the ConvLSTM in this branch projected the initial coordinates maps into hidden states with the same size of the output of branch E.

We used the attention branch to combine the outputs of E and P to construct the spatio-temporal features for the target cell, *P*(*ST*^*α*^|*ST,Q*^*α*^). It fused the populate latent state and target representations by leveraging the attention mechanism. In an attention module (Wang et al., 2017), there are two terms: mask branch and trunk branch. Trunk branch is used for feature extraction in attention module. Mask branch learns the same size feature as trunk branch and behaves as feature selector during forward inference. The output of an attention module is *σ*(*M*(*x*) ∗ *T*(*x*)), where *σ* is a sigmoid function, and *M* and *T* represent the mask branch and trunk branch with input *x*. In the proposed work, the fused deep representation generated from branch E served as the trunk output (Eq. 7), where ⊕ is the concatenate operation on two vectors. And the *Q*^*α*^ was treated as a deep mask in latent space. Because the ConvLSTM applied an equivalent attention learning to each channel of its input, we implement this attention module in two steps. First, we concatenated all deep representation generated from branch E and P as one hidden state tensor (Eq. 8). Then we selected and compressed the spatio-temporal feature specific to the target by applying two ConvLSTMs layers. After features fusion, we injected the *ST*^*α*^ into the task specific networks as the input.

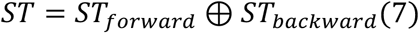

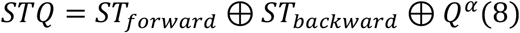

### Task Specific Networks

As mentioned above, the task specific networks can project the latent space into a two-dimensional response map to represent object activities. The task specific networks for tracking task, mitosis detection task, and apoptosis detection task are shown in Figure 2. For each task, we utilized a similar architecture to accomplish the projecting. Given the spatio-temporal feature *ST*^*α*^ for the target cell *α*, the first convolutional layer takes the common input with filter kernels of size 5-by-5 to cover the area around a pixel. The next convolutional layer with filter kernels of size 1-by-1 produced a fusion map, *h*_*task*_, for the corresponding tasks. A batch normalization layer was applied between the convolutional layers. Then, the network outputted the predictive response map by an extra sigmoid layer. We express the output of the model by Eq. 9, where ⊕ is the concatenate operation on two vectors, and *R*_*s*_, *R*_*w*_, *R*_*z*_ are the response map of the cell tracking task, mitosis detection task, and apoptosis detection task, respectively. To convert the response map to the coordinates of the target cell, we selected the *k* sorted local maxima of the response map to indicate the spatial locations. And all local maxima values were needed to be larger than the corresponding task threshold: *T*_*task*_. In this study, we assume that there was up to one peak in a response map, which means *k* = 1.

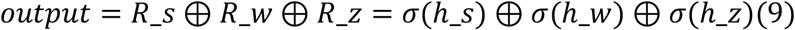

**Figure 2.**
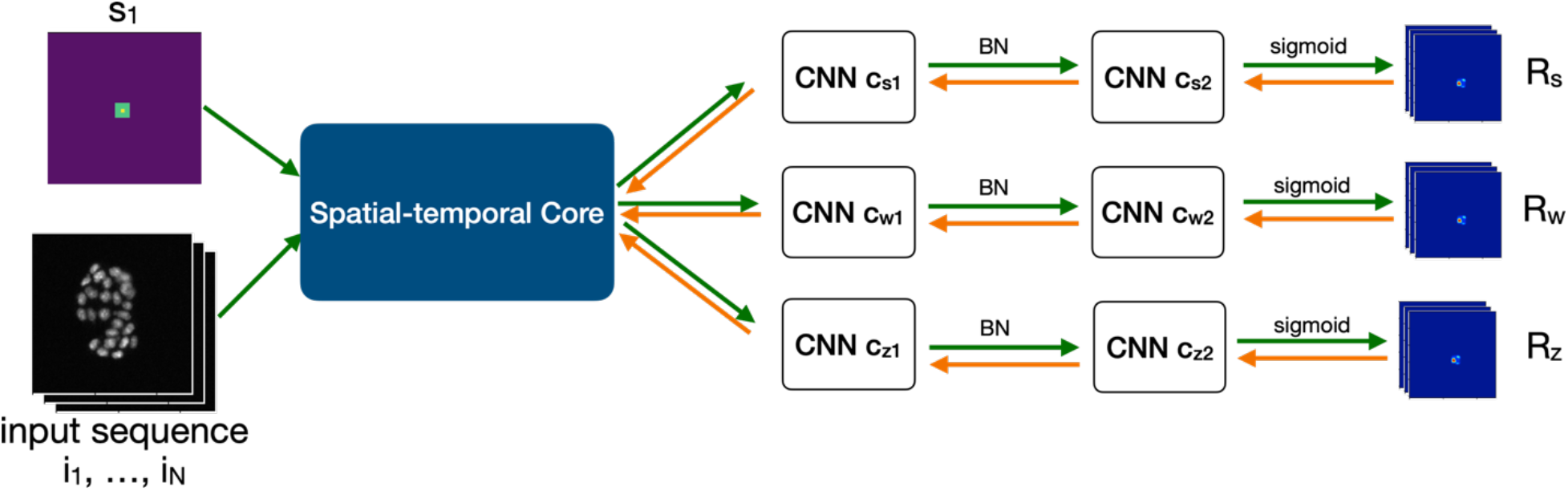
The architecture of the proposed model with three task specific networks for tasks: *P*(*S*^*α*^|*ST*^*α*^) denotes the cell tracking task, *P*(*W*^*α*^|*ST*^*α*^) represents the mitosis detection task, *P*(*Z*^*α*^|*ST*^*α*^) indicates the apoptosis task. The green arrow indicates the inference paths. The orange arrow indicates the backpropagation paths.

### Model Training

In this multi-task training problem, we focused on both cell tracking and biological events detection. We designed a loss function (Eq. 10) for optimization, where *λ*_*s*_, *λ*_*w*_, *andλ*_*z*_ are the inter-task trade-off parameters for tracking, mitosis detection and apoptosis detection; 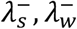, *and*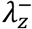 are the inner-task trade-off parameters; *x* and *y* denote the spatial coordinates; *t* indicates the time step. Because the area of background was bigger than the area target cell in ground truth map, the inner-task parameters were defined as 0 < *λ*^−^ ≤ 1.

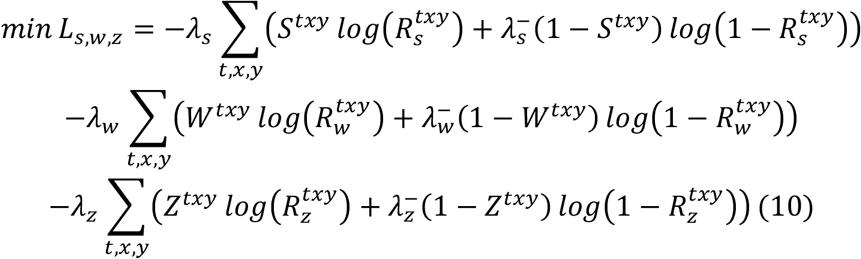

We introduced an incentive for the spatial representation of the coordinates *S*^*α*^ to encourage the model to learn patterns around a cell as well as to learn to focus on the central point of the target cell. For the ground truth map, instead of using only one pixel with value 1 and all others with value 0, a small square of size 3 × 3 and a large square of size 10 × 10 centering at coordinates of the target cell *C*^*α*^ were assigned values of 1 and 0.7, respectively.

In practice, we implemented the proposed model with a TensorFlow framework. We used RMSProp with mini-batch and back-propagation through time as optimization algorithm (Werbos, 1990; Tieleman & Hinton, 2012), with a learning rate of 10^−3^ and a decay rate of 0.9. We initialized the training parameters by a Gaussian distribution with a mean value of 0 and a variance of 0.01. All the parameters in the loss function were differentiable. We set the inter-task trade-off parameters as 1, the inner-task trade-off parameters as 0.7, and the task threshold as 0.5. Each ConvLSTM layer had 32 hidden states. All the input-to-state and state-to-state kernels were 5 × 5. We trained our Cell-STN models over 100 epochs.

### Performance evaluation

In this study, we compared the Cell-STN with two benchmarks, EllipTrack and BaxterAlgorithm algorithms (Magnusson et al., 2015; Tian, Yang & Spencer, 2020). On the HeLa dataset, the parameter values provided with the EllipTrack and BaxterAlgorithm were used. Because of different use cases, these values were fine-tuned on both training set and testing set. For the in-house datasets, we carefully tuned the parameters of the two benchmarks on training set and report the evaluation performance on testing set. Because the benchmarks were trained by manually annotated masks, we did not report the spatial distance and temporal distance for these methods. The benchmarks used the cell nuclear channel (CFP channel in this study) as the main source to track the cells. For fair comparison, we trained and evaluated two versions of the proposed Cell-STN models. The Cell-STN c/3 models were to leverage input images with three-channels. The Cell-STN c/1 models were to take input images only with a single channel (CFP channel).

### Statistical Analysis

We conducted two types of evaluation experiments on the testing sets to show the performance of the Cell-STN. For the cell tracking task, we calculated the peak-wise precision, track-wise precision, and end-peak precision to assess localization ability and tracking capability of the proposed models. The peak-wise precision was the ratio between the number of correct peaks and the total number of the predicted peaks. The track-wise precision was a ratio between the number of correct tracks (all peaks in the track were correct predictions) over the total number of tracks. The end-point precision was defined as the ratio between the number of correct peaks (located in the last frame of each track) over the total number of tracks. Because we split long-period movies into multiple small-piece sequences, it was a practical strategy to track cells over a long time period that using the last predictions of each sequence as the initial location of the following frames. Therefore, the end-peak precision measured the potential capacity to apply this tail-to-nose strategy. A predicted peak was considered as true positive if it is spatially within 10 pixels and temporally within 1 frame from the ground-truth annotation that followed the true position detection definition in previous works (Huh & Chen, 2011; Phan et al., 2019). We also measured the spatial distance between the predicted peaks and the ground truth coordinates. We use the RMSE to compute this distance in the unit of pixels.

For the biological event detection tasks, including mitosis detection and apoptosis detection, we adopt the F-1 score, defined as the harmonic mean between precision and recall, as the detection metric. Since both true positive and true negative predictions were important to the detection tasks, we also reported the model performance in terms of accuracy that is the ratio between true predictions and total predictions. In addition to the spatial distance, we also reported the temporal distance between the predictions and ground truth in the unit of frames.

## Results

### Cell Tracking Task

Figure 3 shows an example that reveals the tracking output of the Cell-STN performance on the HCT116 dataset. Although the moving speed of HCT116 cells is faster than average, the predicted location points are always close to the ground truth. The response map of the first frame shows a compact square with value close to 1, which means the model is extremely confident with this result. This is because the initial location of the target cell is given to the model as an input. As the time increase, the highlighted area becomes disperse since the surrounding cells are considered as potential tracking candidates. However, there is a huge distinction between the true target and surrounding candidate. Furthermore, the peak value is always larger than 0.8. According to our visual inspection of the model outputs, the Cell-STN has the capability to track a cell given the initial location and imaging sequence.

**Figure 3.**
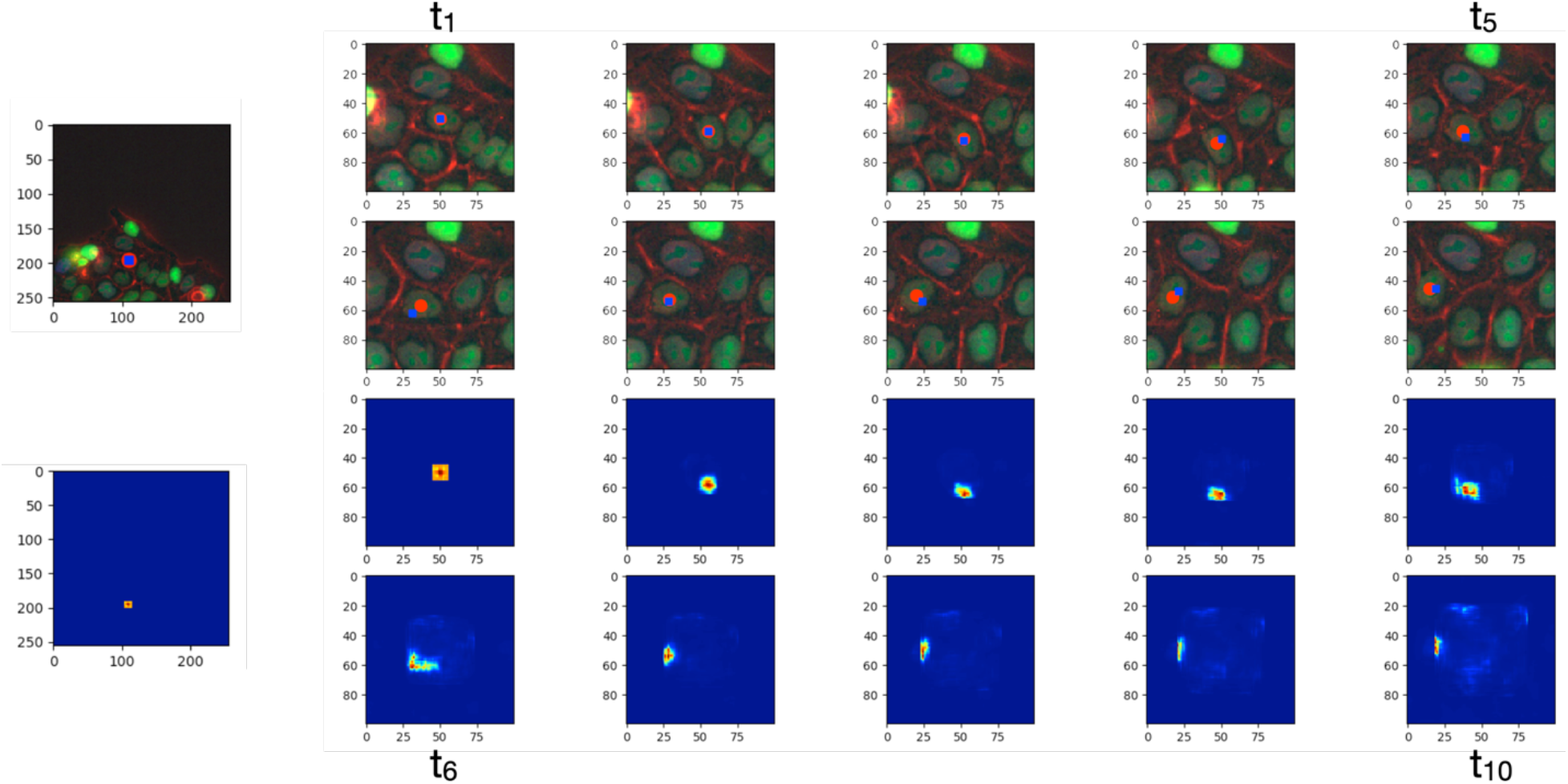
An example of the cell tracking task performed on HCT116 dataset. The first column shows the start position of a target cell. The *t*_1_to *t*_10_ indicate the image frame from 1 to 10 in two rows. In the upper half frames, the red dots indicate the ground truth location of the target cell.

For better illustration, we summarize the evaluation results of models in Table 1. Overall, the proposed Cell-STN has a more powerful ability to track cells in different environments. For the in-house datasets, both versions of the proposed works, three-channels and single-channel models, consistently outperformed the benchmarks in terms of peak-wise precision. Although the benchmarks achieved similar performance in terms of track-wise precision on the U2OS and HCT116 datasets, the Cell-STN kept the superiority on the end-peak precision. All models awarded lower performance on the HCT116 dataset that was expected because the HCT116 dataset was the most challenging dataset for the tracking task in this study. For the public dataset (HeLa), all models had similar performance on the tracking task.

**Table 1.** Cell tracking performance on four datasets.

|  | Peak-wise<br>Precision | Track-wise<br>Precision | End-peak<br>Precision | Spatial<br>Distance |
| --- | --- | --- | --- | --- |
| **MCF7** |  |  |  |  |
| **EllipTrack** | 0.95 | 0.85 | 0.88 | - |
| **BaxterAlgorithm** | 0.96 | **0.88** | 0.90 | - |
| **Cell-STN (c/1)** | **0.97** | 0.87 | 0.95 | 3.03 |
| **Cell-STN (c/3)** | **0.97** | **0.88** | **0.96** | **2.79** |
| **U2OS** |  |  |  |  |
| **EllipTrack** | 0.91 | **0.82** | 0.88 | - |
| **BaxterAlgorithm** | 0.93 | 0.81 | 0.91 | - |
| **Cell-STN (c/1)** | **0.96** | 0.80 | 0.91 | 3.60 |
| **Cell-STN (c/3)** | **0.96** | **0.82** | **0.94** | **3.24** |
| **HCT116** |  |  |  |  |
| **EllipTrack** | 0.81 | 0.73 | 0.76 | - |
| **BaxterAlgorithm** | 0.80 | **0.69** | 0.72 | - |
| **Cell-STN (c/1)** | 0.89 | 0.67 | 0.79 | 5.22 |
| **Cell-STN (c/3)** | **0.90** | **0.69** | **0.82** | **4.77** |
| **HeLa** |  |  |  |  |
| **EllipTrack** | 0.97 | 0.91 | 0.94 | - |
| **BaxterAlgorithm** | **0.98** | **0.92** | **0.97** | - |
| **Cell-STN (c/1)** | **0.98** | **0.92** | **0.97** | **3.24** |

And the blue dots indicate the predicted location of the target cell. The lower half frames show the corresponding response maps displayed as heat maps.

### Mitosis Detection Task

We present an example of the mitosis detection task in Figure 4. In the upper half frames, both the predicted mitosis event and the ground truth are shown in the 7th frame and close to each other spatially. In the lower half frames, the peak with high values only exists in the 7th response map that shows a strong correlation with the ground truth.

**Figure 4.**
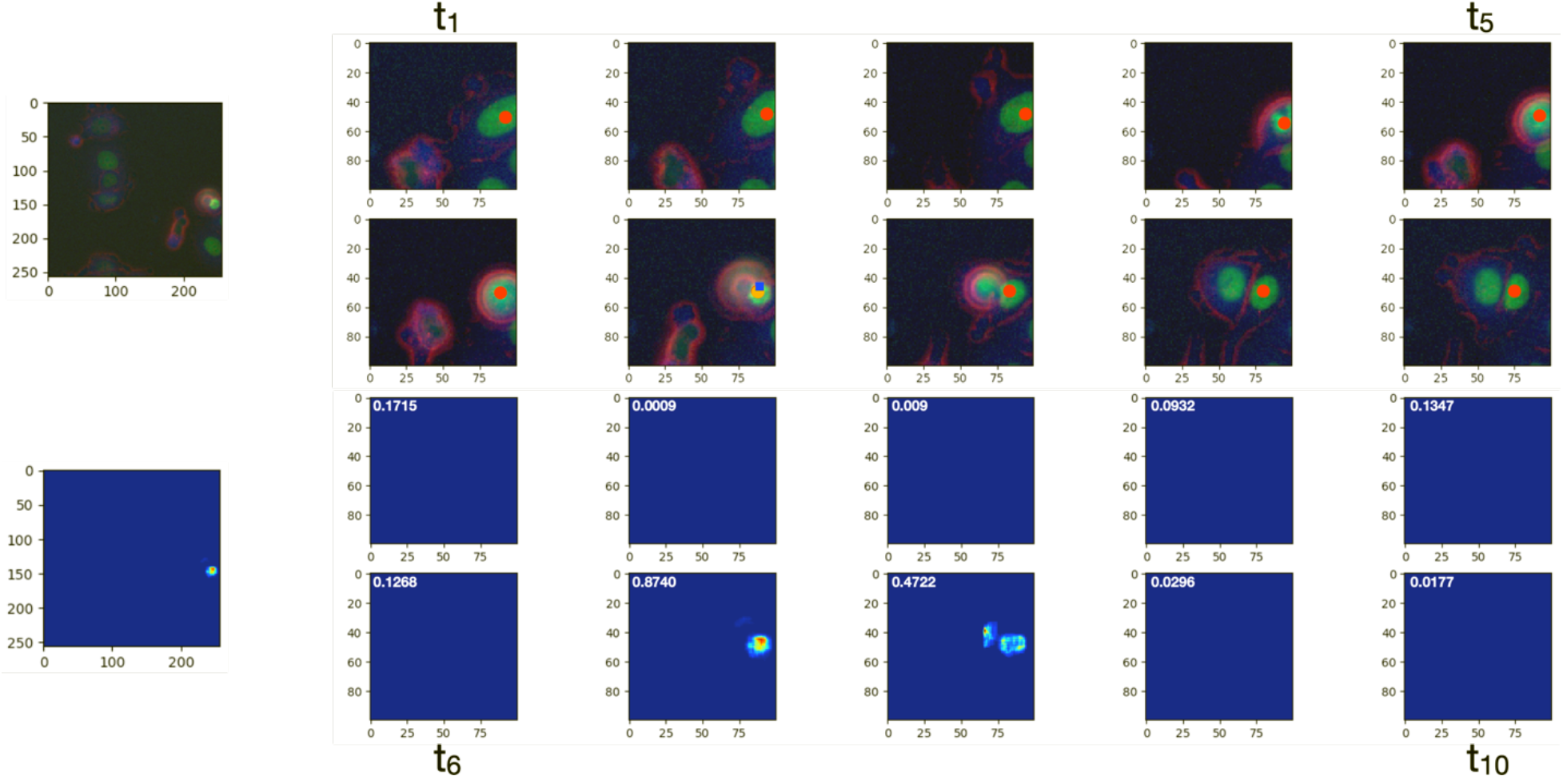
An example of the mitosis detection performed on U2OS dataset. The first column shows the initial position of a target cell. The *t*_1_to *t*_10_ indicate the image frame from 1 to 10 in two rows. In the upper half frames, the red dots indicate the ground truth location of the target cell. And the blue dot indicates the predicted mitosis event of the target cell. The orange dot highlighted the spatial and temporal ground truth of a mitosis event. The lower half frames show the corresponding response maps displayed as heat maps. The mitosis event shows in the 7^th^ frame.

We also verified the effectiveness of the Cell-STN on the mitosis detection task. We list the evaluation results in Table 2. The three-channel Cell-STN achieved the highest performance in terms of accuracy and F1 score across the three in-house datasets. The margin between the single-channel Cell-STN and the benchmarks is also obvious. Additionally, we also calculated the frequency distribution of birth event timing errors of the cell division detection for Cell-STN. The three-channel method achieves higher temporal accuracy than the single-channel approach.

**Table 2.** Mitosis detection performance on four datasets under different temporal thresholds.

|  | Accuracy | F-1 Score | Temporal Distance | Spatial Distance |
| --- | --- | --- | --- | --- |
| **MCF7** |  |  |  |  |
| **EllipTrack** | 0.61 | 0.32 | - | - |
| **BaxterAlgorithm** | 0.50 | 0.39 | - | - |
| **Cell-STN (c/1)** | 0.72 | 0.69 | 1.34 | 9.28 |
| **Cell-STN (c/3)** | **0.82** | **0.79** | **1.13** | **7.51** |
| **U2OS** |  |  |  |  |
| **EllipTrack** | 0.79 | 0.52 | - | - |
| **BaxterAlgorithm** | 0.65 | 0.32 | - | - |
| **Cell-STN (c/1)** | 0.77 | 0.75 | 0.91 | 7.49 |
| **Cell-STN (c/3)** | **0.85** | **0.87** | **0.42** | **4.94** |
| **HCT116** |  |  |  |  |
| **EllipTrack** | 0.48 | 0.28 | - | - |
| **BaxterAlgorithm** | 0.37 | 0.19 | - | - |
| **Cell-STN (c/1)** | 0.83 | 0.78 | 0.68 | 10.19 |
| **Cell-STN (c/3)** | **0.86** | **0.87** | **0.37** | **7.16** |
| **HeLa** |  |  |  |  |
| **EllipTrack** | 0.84 | 0.81 | - | - |
| **BaxterAlgorithm** | 0.82 | 0.80 | - | - |
| **Cell-STN (c/1)** | **0.86** | **0.87** | **0.34** | **4.89** |

### Apoptosis Detection Task

For the apoptosis detection task, we show an example performed on the U2OS dataset in Figure 5. Both the prediction and the apoptosis ground truth are shown in the 9th frame. Although the peaks in the last two response map were larger than the threshold *T*_*apoptosis*_, the model selected the peak in the 9th as the final prediction because this peak had the highest response map value.

**Figure 5.**
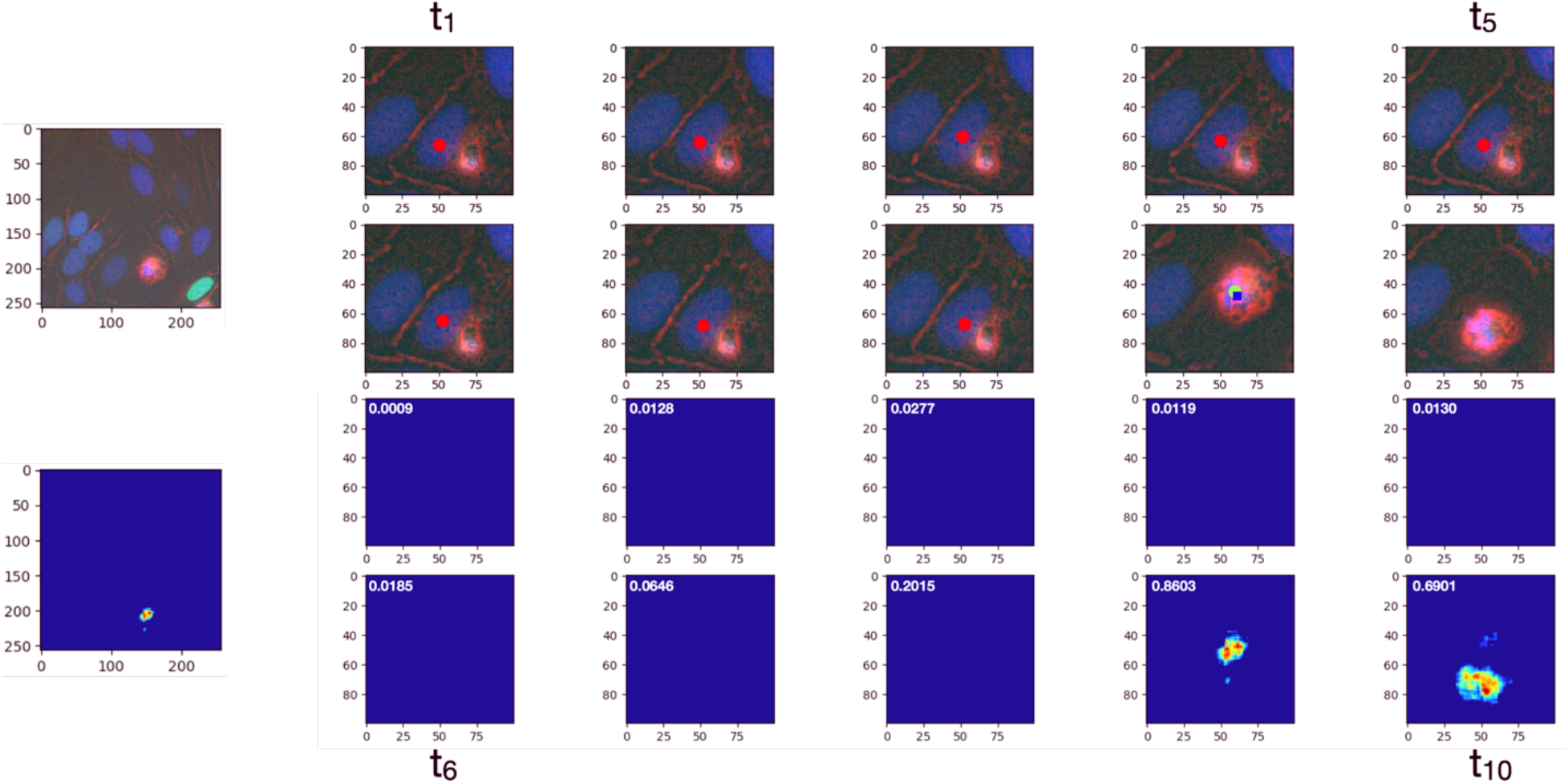
An example of the apoptosis detection performed on U2OS dataset. The first column shows the initial position of a target cell. The *t*_1_to *t*_10_ indicate the image frame from 1 to 10 in two rows. In the upper half frames, the red dots indicate the ground truth location of the target cell. And the blue dot indicates the predicted apoptosis event of the target cell. The green dot highlighted the spatial and temporal ground truth of an apoptosis event. The lower half frames show the corresponding response maps displayed as heat maps. The highest peak value shows in the 9th response map.

In Table 3, we show the comparison of the performance of our Cell-STN models with mask-based benchmarks. Since cell apoptosis events were rare in the HeLa dataset, this dataset was excluded in this evaluation. In terms of accuracy and F-1 score, the three-channel Cell-STN models outperformed other algorithms on the apoptosis detection. Comparing to the benchmarks, the single-channel Cell-STN models achieved better performance on MCF7 and U2OS datasets and similar performance on the HCT116 dataset. Compared to the single-channel Cell-STN, the three-channel version had better performance in terms of the temporal distance and spatial distance.

**Table 3.** Apoptosis detection performance on three datasets under different temporal thresholds.

|  | Accuracy | F-1 Score | Temporal Distance | Spatial Distance |
| --- | --- | --- | --- | --- |
| **MCF7** |  |  |  |  |
| **EllipTrack** | 0.69 | 0.58 | - | - |
| **BaxterAlgorithm** | 0.68 | 0.32 | - | - |
| **Cell-STN (c/1)** | 0.89 | 0.82 | 1.28 | 8.47 |
| **Cell-STN (c/3)** | **0.95** | **0.92** | **0.45** | **4.38** |
| **U2OS** |  |  |  |  |
| **EllipTrack** | 0.57 | 0.46 | - | - |
| **BaxterAlgorithm** | 0.57 | 0.44 | - | - |
| **Cell-STN (c/1)** | 0.75 | 0.69 | 1.58 | 29.58 |
| **Cell-STN (c/3)** | **0.89** | **0.89** | **0.52** | **6.37** |
| **HCT116** |  |  |  |  |
| **EllipTrack** | 0.49 | 0.30 | - | - |
| **BaxterAlgorithm** | 0.51 | 0.35 | - | - |
| **Cell-STN (c/1)** | 0.49 | 0.39 | 1.87 | 11.89 |
| **Cell-STN (c/3)** | **0.79** | **0.77** | **1.51** | **8.34** |

### Ablation Study

To investigate the contribution of the ground truth mask in the proposed Cell-STN, we trained a Cell-STN model using the cell masks as ground truth and conducted a comparison with the original Cell-STN model on the HeLa dataset. Formally, the cell masks are defined as 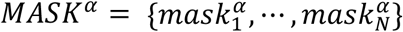 and 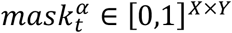. If the location (*x, y*) belongs to the cell *α*, the (*x, y*)^*th*^ entry of the *MASK*^*α*^ equals 1; otherwise, it equals 0. Because of the same variable dimension, we replaced the coordinates maps *S*^*α*^ by *MASK*^*α*^ to form the mask version of the Cell-STN. We leveraged the ground truth masks provided in the HeLa dataset to train the Cell-STN models and reported the comparison results in Table 4. The results showed the original Cell-STN without mask achieved a similar or even better performance on the HeLa dataset.

**Table 4.** Performance comparison between with mask and without mask.

| **Cell Tracking Task** |  |  |  |  |
| --- | --- | --- | --- | --- |
|  | Peak-wise Precision | Track-wise Precision | End-peak Precision | Spatial Distance |
| **Cell-STN (w/o mask)** | **0.98** | **0.92** | **0.97** | **3.24** |
| **Cell-STN (with mask)** | 0.95 | 0.83 | 0.93 | 4.82 |
| **Mitosis Detection Task** |  |  |  |  |
|  | Accuracy | F-1 Score | Temporal Distance | Spatial Distance |
| Cell-STN (w/o mask) | 0.86 | **0.87** | 0.34 | 4.89 |
| Cell-STN (with mask) | **0.87** | 0.84 | **0.36** | **4.67** |

## Discussion

In this study, we developed and evaluated a deep learning approach that can automatically track cells and detect both mitosis and apoptosis events. The model was trained and tested on 3 in-house datasets and 1 public dataset. The proposed Cell-STN consisted of two parts: spatio-temporal core network and a task specified network. Compared to human expert annotated ground truth, the model attained a high level of performance.

Our finding that the proposed models awarded the best performance on cell tracking, mitosis detection, and apoptosis detection is of particular interest. In contrast to the two benchmarks (EllipTrack and BaxterAlgorithm), the proposed Cell-STN showed consistantly better performance on the three tasks. On the cell tracking task, although the benchmarks achieved similar performance in terms of track-wise precision on the U2OS and HCT116 datasets, the Cell-STN kept the superiority on the end-peak precision. This showed the effectiveness of the use of ConvLSTM to makes predictions based on both short-term appearance and long-term dependencies. As a result, the Cell-STN had the capability to self-correct the following tracking predictions even if one of the previous predictions was wrong. On the mitosis detection task, the Cell-STN showed a greater advantage than the two benchmarks. It is worth noting that we did not guide the Cell-STN models to learn the features of daughter cells in training. Instead, in the training process, we only used the binary label of the presence of the mitosis events with a coordinate map to train the Cell-STN models. In Figure 4, the model highlights both daughter cells in the following frame after mitosis occurred. This shows the model’s strong capacity to learn useful features from implicit cues. And the Cell-STN has the potential to be adopted in a weakly supervised learning manner. For the apoptosis detection task, the Cell-STN also showed advantages. In our annotating process, experimentalists stopped to track a cell immediately after apoptosis. Consequently, no ground truth was provided to the model after apoptosis during the training process. However, the model generated response map tended to highlight the dead cell for a short period because cells typically showed cohesive and last few frames after they die. In the next few frames, the boundary of these dead cells became blurred, and the brightness decreases until they are no longer visible. Therefore, the Cell-STN has not only better performance, but also more useful functions than the two benchmarks.

Additional spatio-temporal information is useful for performance improvement. In this study, another interesting finding is that the performance becomes better when we increased the number of input channels. For example, the three-channel Cell-STN achieved higher end-peak precision and spatial distance on cell tracking task and temporal accuracy on the mitosis detection task. The three-channel models also awarded better performance on the apoptosis detection. We believed the additional information (e.g., nuclei information) possibly makes the cell tracking more accurate. This finding is constant with previous studies. The DNNs model have a higher capacity than the performance of deep learning algorithms can be improved by providing more data and information (Esteva et al., 2019). Additionally, the DNNs algorithm can directly learn features from raw input and discover useful and unrecognized patterns in high-dimensional data (Lipton et al., 2015). Therefore, providing more spatio-temporal cues can improve the overall model performance.

Interesting, we also found the extremely accurate mask labels cannot improve the model performance and the coarse mask labels can improve model robustness. The original Cell-STN models trained by coarse cell masks achieved a similar or even better performance than the Cell-STN models trained by accurate cell mask on the HeLa dataset. In training, because the coordinates map *S*^*α*^ was set to be a two-level fix-size square to approximate the spatial location of the center of a cell, it can be seen as an extremely coarse cell mask. In inference, the model selected the local maxima of the response blobs that filtered out the noisy information regarding the shape of cells. The coarse mask in training and feature selection in inference formed a bottleneck that increased the robustness of models. On the other hand, although the accurate cell mask provided additional information, they mainly focused on the size and shape of the cells that barely contributed to the final inference process. In addition, the accurate cell masks treat all positions within the cell with equal importance, which generates confusion to the bottleneck feature selection process. Therefore, we considered that the accurate mask-level labels have a negative impact to the optimization of Cell-STN models.

There are several limitations to our study. First, the sizes of datasets were imbalanced. The evaluation results on small datasets may be not as powerful as the evaluation results on big datasets. Second, on the HeLa dataset, the parameter values provided with the EllipTrack and BaxterAlgorithm were fine-tuned on both training set and testing set. Therefore, the performance of the two algorithms may be over-estimated. Third, we acknowledge that the ablation study was preliminary as we only conducted it on one dataset. Further study is required to include more datasets to investigate the impact of the mask ground truth.

Our study has several strengths. Most importantly, the proposed Cell-STN showed the better performance and more advantages than the two benchmarks, EllipTrack and BaxterAlgorithm, on four datasets. Additionally, we applied weakly supervised methods to train Cell-STN models and evaluated the performance of Cell-STN models on real-world datasets. The proposed Cell-STN has higher potential to be used in daily biology research.

## Conclusions

In this study, we presented a new DNNs approach for cell lineage analysis in microscopy images. Compared to the existing state-of-the-art, the proposed work only required coarse cell masks for the network training that avoided the time-consuming step to annotate accurate cell masks. Specifically, we designed the novel Cell-STN, including a spatio-temporal core network for shared cell features regarding appearance and activities and the task-specific networks for the tracking, mitosis detection, and apoptosis detection tasks. We also proposed an innovative loss function and optimization strategy to train the model in a multi-task manner. Finally, we conducted a comprehensive set of experiments on three collected in-house datasets and one public dataset. The comparison results demonstrated that the Cell-STN outperforms other benchmarks on the task of tracking, mitosis detection, and apoptosis detection. We also analyzed the Cell-STN performance under different number of input channel and investigated the impact of the accurate mask-level ground truth. In the future, we will explore the potential of using Cell-STN to predict high-level features using low-level guidance in a weakly supervised learning manner. We will also continue to investigate the impact of accurate mask-level ground truth using larger scale datasets.

## Notes

### Competing Interest Statement

This work was supported by the National Institutes of Health (1R01GM130864-01), the CMMBS training grant (NIH Grant GM132008), and the MCB department training grant (T32-GM008659).

