## Supplemental for "Spatio-temporal feature based deep neural network for cell lineage analysis in microscopy images"

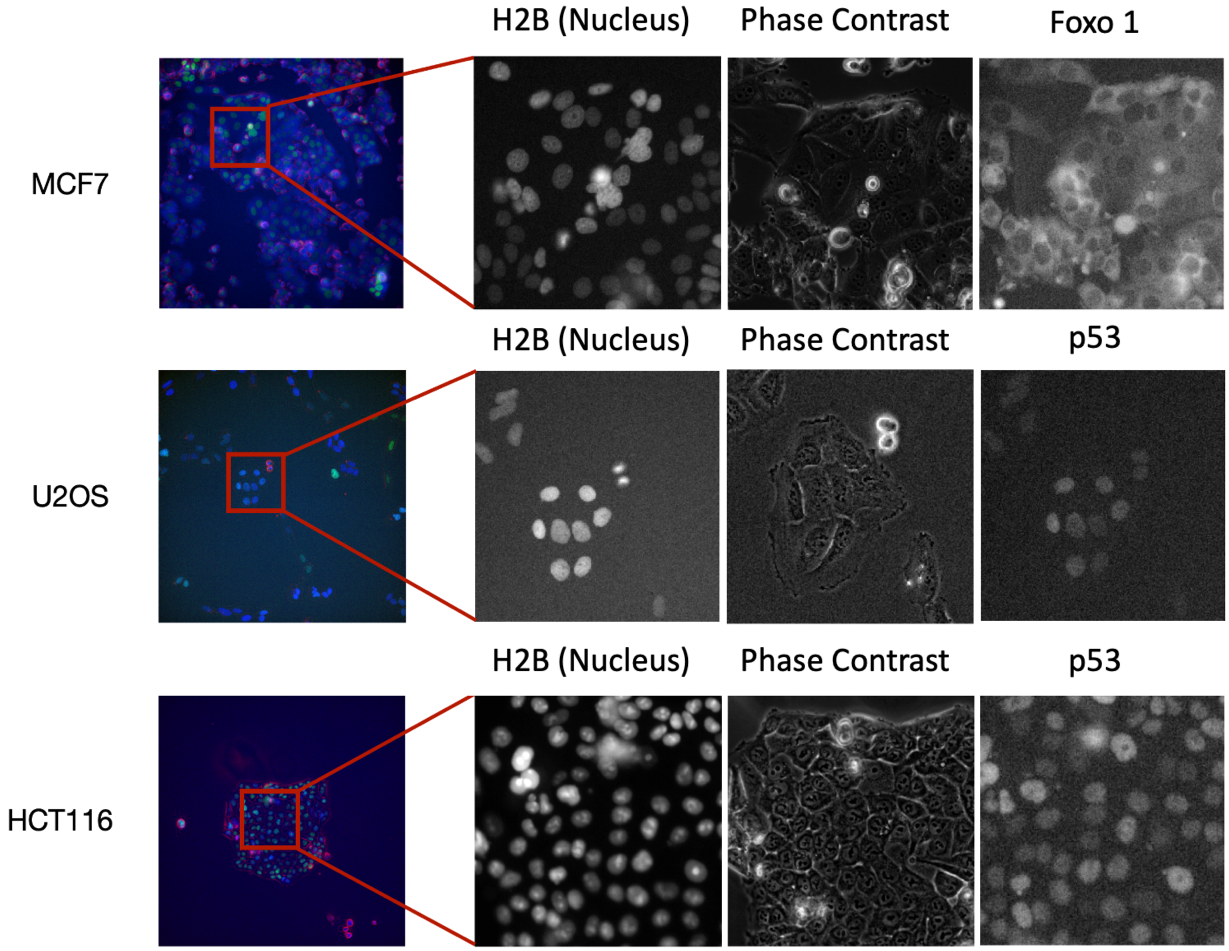


Figure S1. An example of in-house dataset. The first column displays the 3-channel of images in the RGB mode. The CFP channel captured the nucleus information indicated by H2B protein; the bright-field channel captured phase-contrast images; the YFP channel was used for biomarker p53 in the HCT116 and U2OS data sets, and Foxo1 in the MCF7 dataset. The right three columns show the details of the corresponding images within different channels. The images in the same row indicate the same location at a time point.

**Long Short-Term Memory**

The long-short term memory (LSTM), one of the variants of RNN, was originally proposed by Hochreiter and Schmidhuber (Hochreiter & Schmidhuber, 1997). It consists of different memory cells and gating components to store information and control the flow of the content being stored. In this study, we used the formulation of LSTM reported by Graves (Graves, 2013). Figure S2 shows an LSTM unit with a memory cell, holding a state $c_t$ at time $t$, which acts as an accumulator of the state information. Three self-parameterized controlling sigmoidal gates, including an input gate $i_t$, a forget gate $f_t$, and an output gate $o_t$, govern the access to this memory cell as shown in Figure S2. For every new frame, two external sources and an internal source are received at the four terminals of a LSTM unit. The external sources include the input frame $x_t$ and the previous hidden states of LSTM units at the same layer $h_{t-1}$, while the internal source is the memory cell state $c_{t-1}$. The memory cell will accumulate new information if the input gate is activated. If the forget gate $f_t$ is triggered, the LSTM unit will remove the past memory cell status $c_{t-1}$. The output gate $o_t$ controls whether the state $c_t$ will be propagated to the final hidden state $h_t$. The gates are activated by passing the accumulated inputs, along with a bias, through a sigmoid function. The memory cell state is computed using the previous memory cell state $c_{t-1}$ after passing through the forget gate, and the inputs at the input terminal after getting through the $tanh$ non-linearity and input gate. The updated memory cell state is then passed through $tanh$ non-linearity and the output gate to get the final output, the hidden state $h_t$. This gating mechanism allows for the back-propagated gradient to be trapped in the memory cell that prevents the vanishing and exploding problem in RNN training. The LSTM unit can be represented by Eq. 1 – Eq. 5, where ($\circ$) denotes the Hadamard operator.

$$i_t = \sigma(W_{xi}x_t + W_{hi}h_{t-1} + W_{ci} \circ c_{t-1} + b_i) \quad (1)$$

$$f_t = \sigma\left(W_{xf} * x_t + W_{hf} * h_{t-1} + W_{cf} \circ c_{t-1} + b_f\right) \quad (2)$$

$$c_t = f_t \circ c_{t-1} + i_t \circ tanh(W_{xc} * x_t + W_{hc} * h_{t-1} + b_c) \quad (3)$$

$$o_t = \sigma(W_{xo}x_t + W_{ho}h_{t-1} + W_{co} \circ c_{t-1} + b_o) \quad (4)$$

$$h_t = o_t \circ tanh(c_t) \quad (5)$$

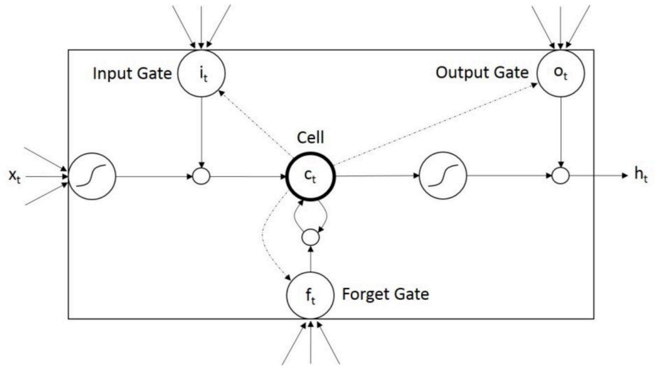


Figure S2. The structure of a LSTM unit

**Convolutional Long Short-Term Memory**

The spatial information is essential in modeling the regularities in image sequences. In this study, we chose the convolutional LSTM (ConvLSTM) as a core block to develop the Cell-STN. The ConvLSTM unit is represented by Eq. 6 -Eq.10. The ConvLSTM preserves spatial and temporal information for modeling long-range dependencies and encodes the spatio-temporal information of cell image sequences (Shi et al., 2015). The major advance in ConvLSTM is the use of 3D tensors for all inputs $x_t$, cell outputs $c_t$, hidden states $h_t$ and gates $i_t, f_t, and\ o_t$ in which the last two dimensions encode the spatial dimensions. The Hadamard products between the weights and inputs $x_t$ or hidden states $h_t$ are then replaced with convolution operators, where $(\circ)$ denotes the Hadamard operator and $(*)$ denotes the convolution operator. As the convolution operation is performed on the inputs and states, padding is required to ensure that the states have the same dimensions as the inputs.

$$i_t = \sigma(W_{xi} * x_t + W_{hi} * h_{t-1} + W_{ci} * c_{t-1} + b_i) \quad (6)$$

$$f_t = \sigma\left(W_{xf} * x_t + W_{hf} * h_{t-1} + W_{cf} * c_{t-1} + b_f\right) \quad (7)$$

$$c_t = f_t \circ c_{t-1} + i_t \circ tanh(W_{xc} * x_t + W_{hc} * h_{t-1} + b_c) \quad (8)$$

$$o_t = \sigma(W_{xo} * x_t + W_{ho} * h_{t-1} + W_{co} * c_{t-1} + b_o) \quad (9)$$

$$h_t = o_t \circ tanh(c_t) \quad (10)$$
